# Rapid Identification and Validation of Novel Rheumatoid Arthritis Drug Treatments using an Integrative Bioinformatics Platform

**DOI:** 10.1101/243998

**Authors:** Aaron C. Daugherty, Carl Farrington, Isaac Hakim, Sana Mujahid, Elizabeth S. Noblin, Andrew M. Radin, Mei-Sze Chua, Mark Rabe, Guy Fernald, Daniel Ford, Marina Sirota, Laura Schaevitz, Andrew A. Radin

## Abstract

The majority of drugs currently used to treat rheumatoid arthritis (RA) act on a small number of immunomodulatory targets. We applied an integrative biomedical-informatics-based approach and *in vivo* testing to identify new drug candidates and potential therapeutic targets that could form the basis for future drug development in RA. A computational model of RA was constructed by integrating patient gene expression data, molecular interactions, and clinical drug-disease associations. Drug candidates were scored based on their predicted efficacy across these data types. Ten high-scoring candidates were subsequently screened in a collagen-induced arthritis model of RA. Treatment with exenatide, olopatadine, and TXR-112 significantly improved multiple preclinical endpoints, including animal mobility which was measured using a novel digital platform. These three drug candidates do not act on common RA therapeutic targets; however, links between known candidate pharmacology and pathological processes involved in RA suggest hypothetical mechanisms contributing to the observed efficacy.

## Main text

Over 20 million people globally suffer from rheumatoid arthritis (RA), an autoimmune disease that leads to inflammation, destruction of bone and cartilage, and joint deformity^1^. This disease is a significant economic burden given that lost productivity and treatment costs approach $46 billion annually in the US. RA drives an $11 billion therapy market that is projected to grow to upwards of $20 billion by 2020^2^.

Despite recent advances in treatment options, disease progression and symptoms are inadequately controlled in 30–50% of RA patients^3^. A deeper understanding of the immune system in RA has led to improved treatment options such as Disease-Modifying Anti-Rheumatic Drugs (DMARDs) that can delay disease progression. However, these therapies have highly variable efficacy and tolerability among patients, and can leave them vulnerable to life-threatening infections due to broad dampening of factors involved in the immune response^4–6^.

The current RA drug development pipeline is heavily enriched for known immunomodulatory and anti-inflammatory targets. Indeed, data from ClinicalTrials.gov revealed that over 80% of all candidates in Phase 2 or Phase 3 clinical trials act through an immune or inflammation target (Table S1). However, such therapies are reported to be ineffective at reducing RA symptoms or slowing disease progression in some patients and have many undesired side effects^3^. Consequently, patients remain in need of selective, efficacious, tolerable, and safe RA treatments.

By using existing Food and Drug Administration (FDA)-approved drugs for new indications, drug repositioning can enable therapies to go directly into preclinical testing and clinical trials, which helps to reduce the overall costs and risks associated with drug development^7^. A well-known example of drug repositioning is sildenafil, which was initially developed to treat angina, and is now indicated for erectile dysfunction^8^ and pulmonary arterial hypertension^9^. Traditional drug repurposing also led to the use of methotrexate, which was originally developed for treatment of cancer, as a first-line therapy for rheumatoid arthritis^10^.

In recent years, a variety of innovative computational methods have been developed to reposition FDA-approved drugs^11^. Such unbiased approaches have identified genes that are targets of approved RA therapies^12^, and could aid in repurposing additional drugs for RA. This strategy has been successful in identifying drug candidates for diseases such as Charcot-Marie-Tooth type 1A^13,14^, which are in Phase III clinical trials. Similar efforts are currently underway for other diseases including oncology and central nervous system disorders^15^.

We developed an integrative biomedical-informatics-based drug discovery platform that aids in drug discovery by leveraging large amounts of biological, chemical, and clinical data in an unbiased manner. Using this platform, we processed data associated with over 22,000 existing drugs, with the goal of identifying drug candidates for repurposing in RA. In this study, we describe drug candidates that appear to target novel pathways for the treatment of RA while maintaining well-characterized toxicology and tolerability profiles. Based on promising efficacy data obtained from preclinical *in vivo* studies, we believe that these candidates have therapeutic potential for RA.

## Results

### Integrative biomedical-informatics-based drug predictions

Most current RA treatments target similar pathways and at least 30% of patients are not adequately treated by these drugs^3,16^. Thus, we set out to predict novel RA treatments acting through mechanisms of action that differ from existing therapies. To do so, we used our DUMA™ Drug Discovery platform, an integrative biomedical-informatics-based software that predicts drug candidates by integrating disparate biological, chemical, and clinical data. In the first stage, multiple large-scale data sets were sourced and loaded into the platform. These datasets included clinical gene expression data, protein interaction networks, drug-protein binding databases, molecular drug structures, and clinical observation data (Fig. 1). During the sourcing and loading process, datasets were carefully analyzed and run through quality control methods to ensure robust and informative data (see Methods).

**Fig. 1.**
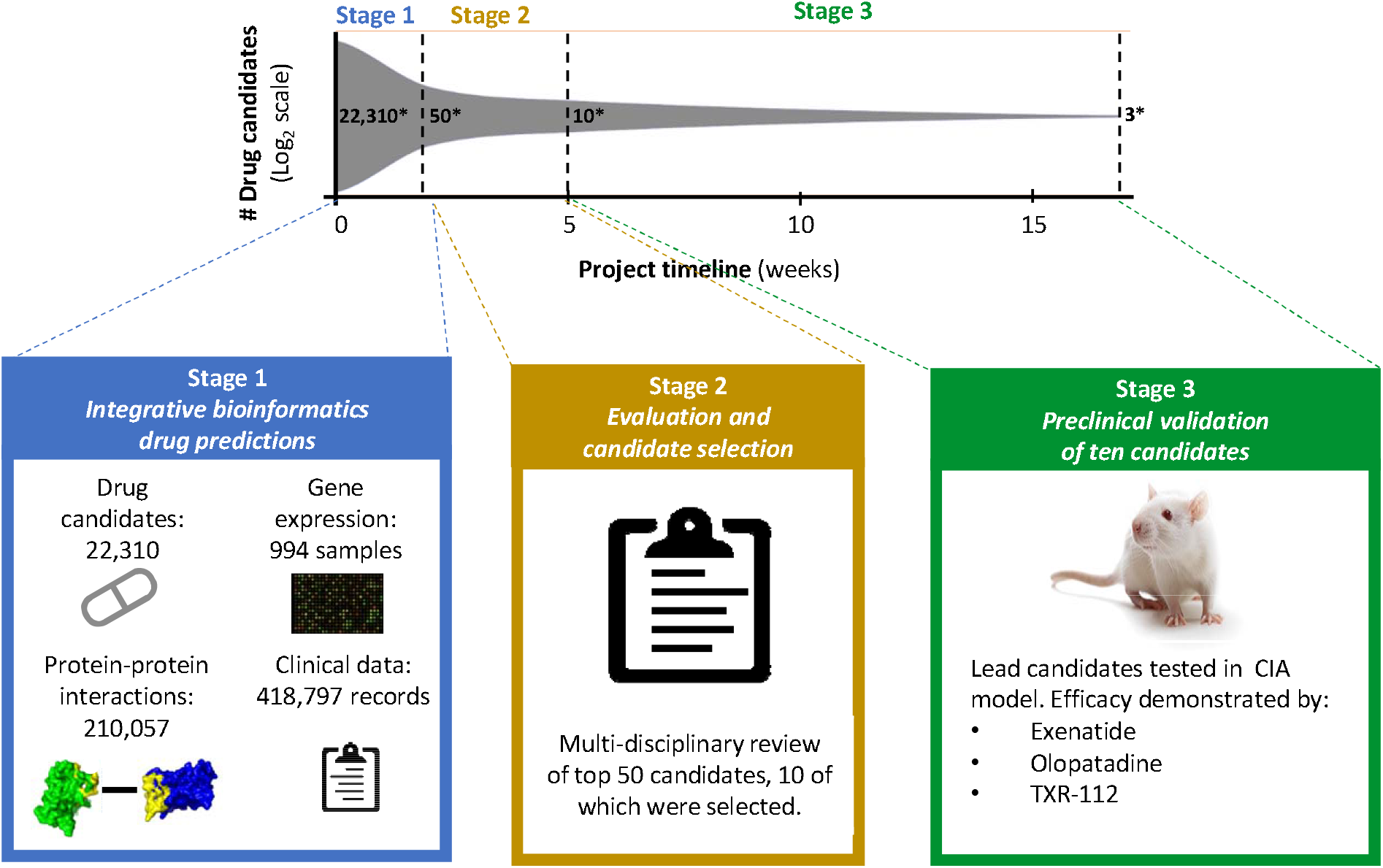
Rapid discovery and *in vivo*-validation of candidates. This integrative biomedical informatics drug prediction approach identifies high potential candidates by integrating diverse data from multiple biomedical sources. In Phase 1, 22,310 large and small molecules present in DrugBank and the Therapeutic Targets Database were scored based on predicted therapeutic potential to yield 50 high-probability candidates. Data used to support drug candidate prediction included: drug-protein interactions, gene expression data from RA patients and healthy controls, protein-protein interaction networks, and records of drug use among patients with and without RA. In Phase 2, algorithm evaluation and candidate due diligence identified ten optimal and novel candidates. In Phase 3, these ten candidates were tested in an *in vivo* model of RA and three lead candidates were identified. The entire project spanned a four-month period. *number of drug candidates.

### Gene expression meta-analysis

A meta-analysis of gene expression changes in RA patients compared to healthy controls was performed using nearly one thousand samples spanning 9 datasets, and 7 tissue and cell types implicated in the pathology of RA (see Methods). For initial verification of this meta-analysis, we performed a gene set enrichment analysis using Reactome, a curated pathway database (Table S2)^17^. A total of 52 and 71 pathways were found to be significantly down-regulated and up-regulated, respectively (Fig. 2; FDR < 0.05). To aid interpretation, the most significant pathways were aggregated into general themes (Fig. 2). Significantly down-regulated groups included several pathways previously associated with RA, including translational and transcriptional processes^18^, phosphatidylinositol metabolism^19^, as well as FGFR and PI3K signal transduction pathways^20,21^. Those pathways significantly up-regulated included expected themes such as immune processes^22^, ECM organization^23^, hemostasis^22^, glycosaminoglycan metabolism^24^, as well as pathways not previously implicated in RA pathogenesis such as insulin signaling. These results indicate that in addition to capturing well established pathways associated with RA, our meta-analysis may be uncovering previously unknown mechanisms involved in the development and progression of the disease.

**Fig. 2.**
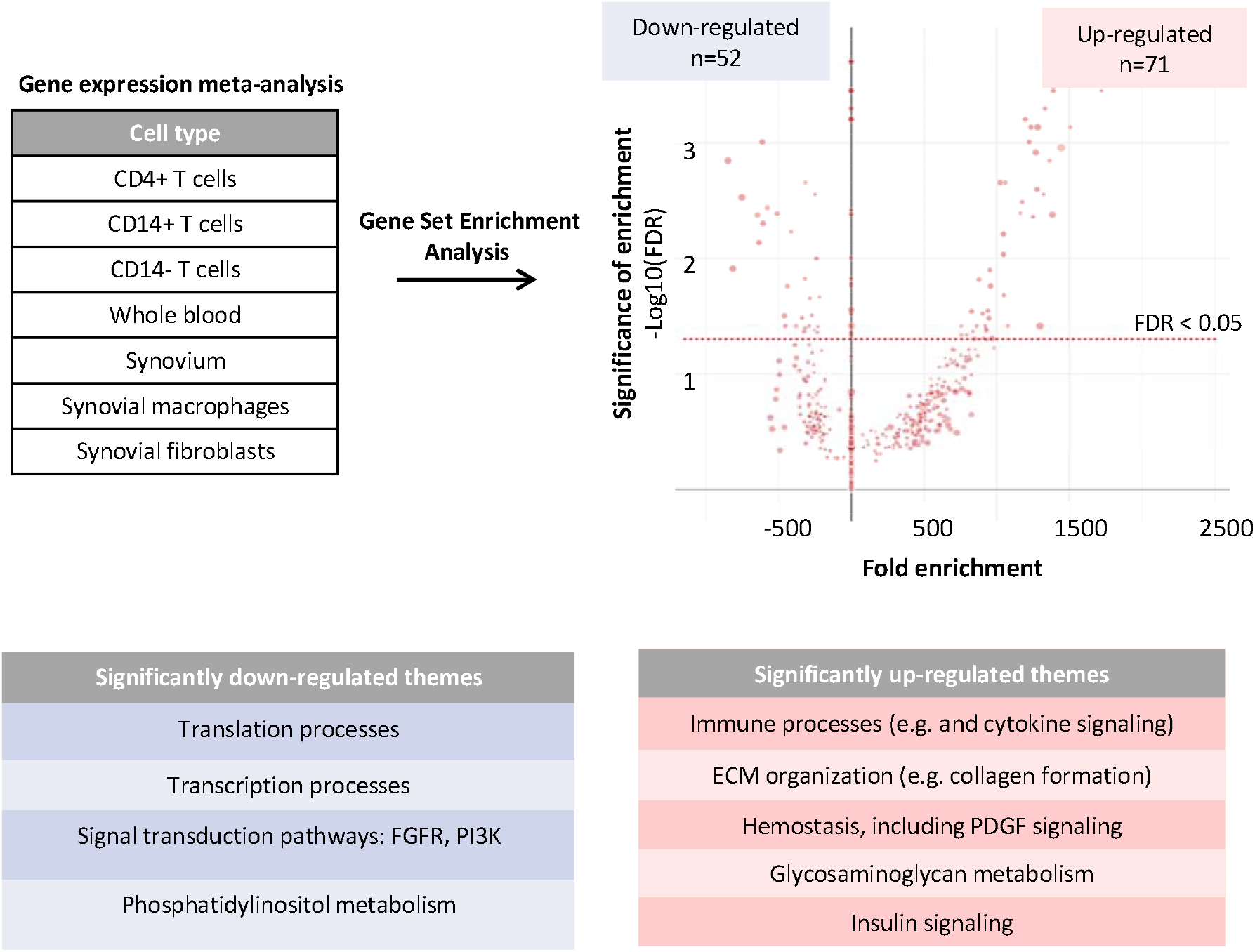
Gene expression data analysis for RA. Publicly available gene expression data for RA patients and healthy controls was collected and analyzed. The graph denotes all pathways that were enriched for down-regulated or up-regulated genes in RA using a gene set enrichment analysis approach. The size of the dot corresponds to the number of all proteins within that pathway that were detected in the RA meta-analysis. The most significant pathways (FDR<0.05) were aggregated into general themes reported in the accompanying tables.

### Drug-protein interaction network

To harness the agnostic and predictive capacity of our gene expression meta-analysis, we employed a systems biology approach. Specifically, information on drug-protein interaction was collected for 22,310 drug candidates from DrugBank^25^ and the Therapeutic Target Database (TTD)^26^ along with protein-protein interaction data from Dr. PIAS^27^. This systems interaction data was integrated with expression changes from the gene expression analysis described above (Fig. 2), and drug candidates were scored according to the number and confidence of their interactions with proteins whose corresponding genes were misexpressed (Fig. 3A). In this method, scoring favors drugs that interact with proteins that are over- or under-expressed in multiple gene expression datasets. The predictive capacity of this scoring method was evaluated by examination of scores for existing RA treatments. This analysis revealed that existing RA therapies, such as adalimumab, were highly enriched among top-scoring candidates (Fig. 3A, p = 1.7×10^−15^). These results demonstrated that by combining differential gene expression and systems biology interactions in a network-based scoring method, existing RA treatments could be re-discovered. This enrichment of existing treatments served as a positive control and provided support to the predictions made using this model.

**Fig. 3.**
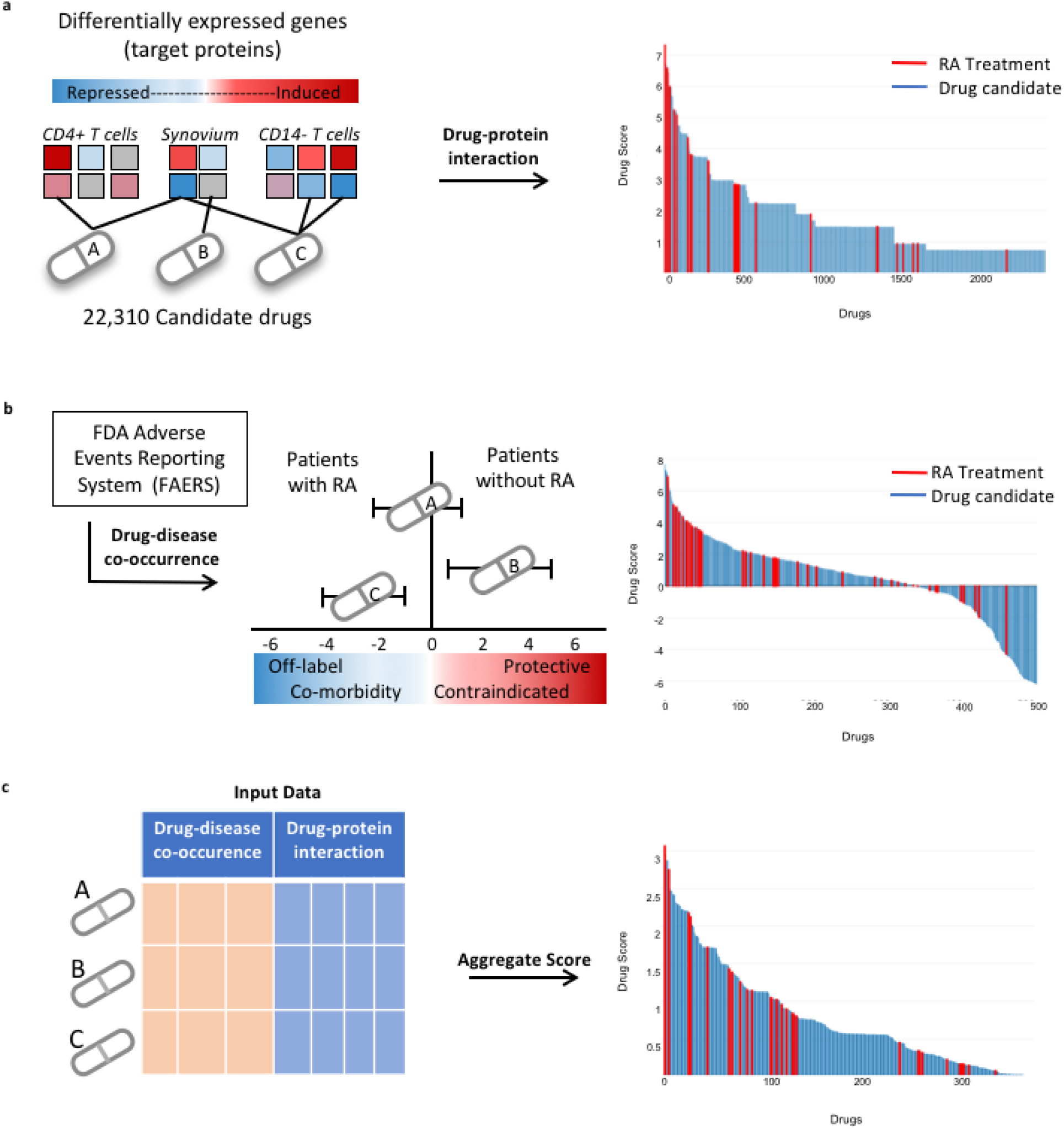
Repositioned drug predictions for RA. (**a**) Information on drug-protein interaction was collected for 22,310 drug candidates. Interaction information was integrated with protein expression changes and candidates were scored according to the number and confidence of interactions. Existing treatments (red bars) for RA were enriched among top-scoring candidates in the drug-protein interaction (p = 1.7×10^-15^). (**b**) Drug-disease co-occurrence scores were calculated by accessing disease diagnoses and medication use from participants in the FDA Adverse Event Reporting System. Existing treatments for RA were enriched among top-scoring candidates (p = 4×10^-3^). (**c**) Scores from B and C were aggregated along with drug characteristics from DrugBank and Therapeutic Target Database into a final score using a machine learning-based approach. RA treatments were most highly enriched in the aggregate scoring method (p = 9×10^-20^).

### Drug-disease co-occurrence

To identify novel RA treatments, we next turned to clinical data sources as an orthogonal approach to the systems biology approach described in Figure 2 and 3a. To do so, drug-disease co-occurrence scores were calculated by accessing disease diagnoses and medication use information from participants in the FDA Adverse Event Reporting System^28^. Drug candidates that tend to co-occur with RA diagnoses, as indicated by positive scores, could represent efficacious off-label treatments or medications used to treat co-morbidities of RA (Fig. 3b). Candidates that tend to co-occur with an RA diagnosis less often than expected by chance, as indicated by negative scores, could represent treatments that are protective against RA (thereby keeping patients from developing RA) or contraindicated in patients with RA (Fig. 3b). For this analysis, we also verified that we were re-discovering existing treatments for RA. Indeed, existing RA treatments were enriched among top-scoring candidates using this drug-disease co-occurrence method (Fig. 3b, p = 4×10^−3^).

### Aggregate scoring

Next, predictions were synergistically integrated from our orthogonal approaches to produce an aggregate score for each drug. Using a proprietary machine learning-based approach, candidate drug efficacy scores and features from the systems biology interaction network and drug-disease co-occurrence were aggregated into a single score (Fig. 3c). As done with the individual scores, the predictive capacity of this final step was evaluated based on the ability to assign high efficacy scores to treatments currently used for RA. Integrating predictions from orthogonal approaches resulted in even greater enrichment of RA treatments (Table S3, Fig. 3c, p = 9×10^−20^), and in fact, among the top 400 overall efficacy-ranked drugs, 53 out of the 55 FDA-approved or clinically used treatments for RA were identified. Using a bootstrapping approach to generate a null distribution, we demonstrated that this enrichment is approximately 12-fold over what might be expected in a disease model with random predictive capacity. This result demonstrated the power of an integrative approach and the robustness of our platform in predicting drug candidates to treat RA.

### In vivo validation of candidates

After establishing credibility of our drug predictions using existing treatments as a benchmark (Fig. 3c), we next sought to identify novel RA drug candidates using a similar approach. To ensure the novelty of these candidates, any existing RA treatments were not considered and we employed a multi-disciplinary team review of our top 50 candidates based on set criteria. During this systematic review process, we excluded any candidates that were previously suggested as RA treatments, or that acted through targets of existing or hypothesized RA treatments as evidenced in the research literature or patents. In addition, any candidates deemed unsafe for RA due to toxicity concerns or were unavailable for commercial purchase were excluded.

Following our systematic review, ten of the 50 highest-ranked candidates by our platform were tested in the collagen-induced arthritis (CIA) model using a novel digital platform. In this standard preclinical model for RA, rats develop autoimmunity against collagen, leading to joint inflammation and degeneration^29^. At Day 0, female Lewis rats were injected with type II collagen in incomplete Freud’s adjuvant to induce arthritis, and a booster was administered on Day 7 (Fig. 4a). Animals were then randomly assigned to treatment groups (8 animals per group). Vehicle or test drug was administered daily starting at Day 9. Dexamethasone (75 μg/kg), a glucocorticoid used to treat autoimmune and inflammatory processes such as RA^30^, was used as an additional control^31^. An efficacious dose of dexamethasone was chosen for these animal experiments^32^. However, this drug is only administered at low doses clinically due to toxicities associated with their chronic use ^33,34^. Therefore, it is highly unlikely for a test drug to outperform dexamethasone in the preclinical setting^35^.

**Fig. 4.**
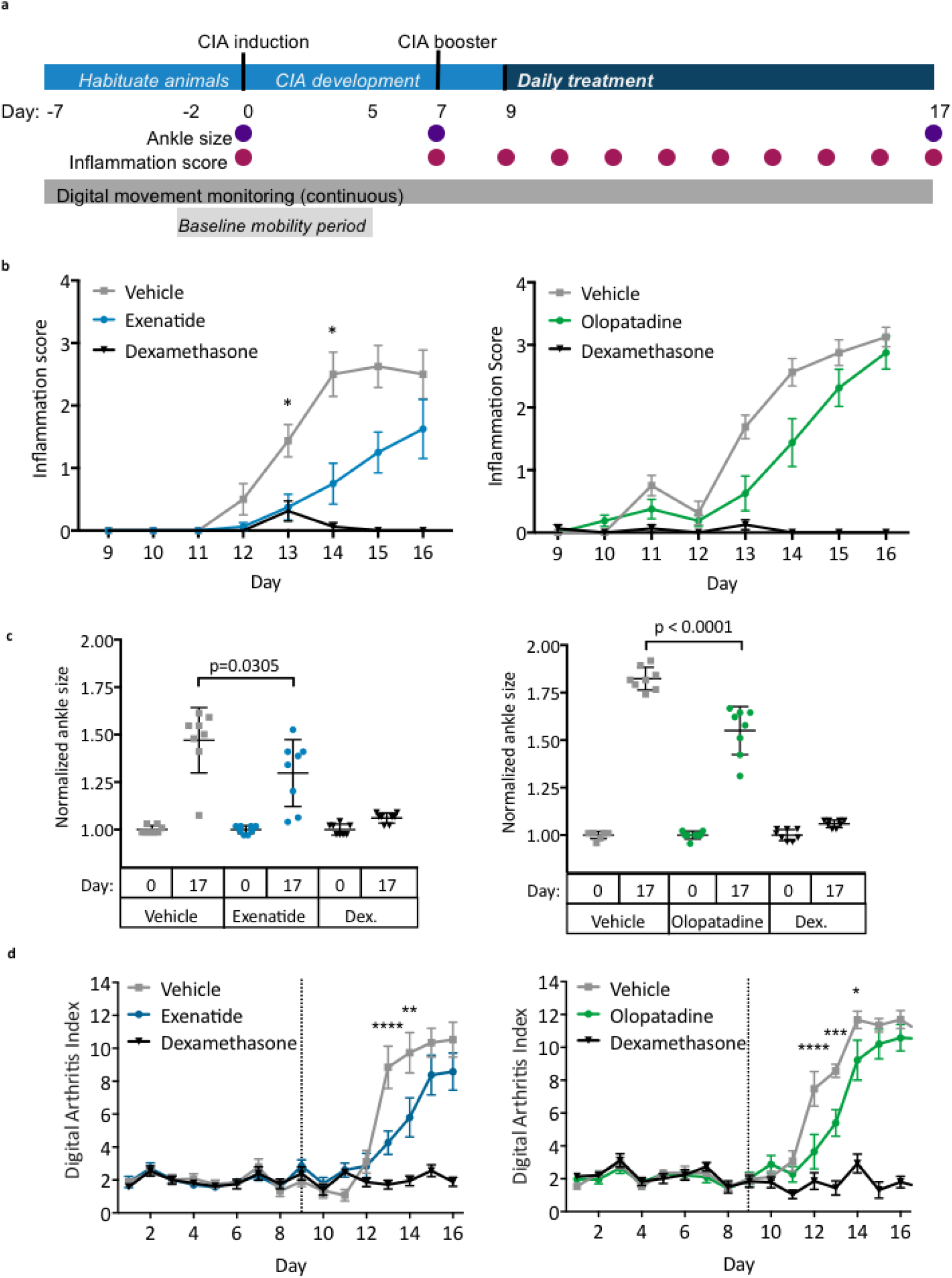
Preclinical validation of candidates. (**a**) Schematic overview of the study design and endpoint measurements. (**b**) Hind-limb inflammation were scored daily and scores for left and right limbs for each animal were averaged in each group. (**c**) Sizes of left and right ankles for each animal were summed, then normalized to ankle size at Day 0 in each group. (**d**) Digital Arthritis Index was calculated using continuous digital monitoring of animal movement. Higher Digital Arthritis Index scores correspond to more severely impaired mobility. Data graphs represent the mean and SEM (n=8 per group). Dex.: dexamethasone; *p < 0.05; ** p < 0.01; *** p < 0.001;**** p < 0.0001, two-way ANOVA with Tukey’s multiple comparison test (ankle size, digital arthritis index) or Kruskall-Wallis with Dunn’s correction (inflammation scores).

Multiple endpoints were measured to assess disease progression. These included daily scoring of hind-limb inflammation and ankle width measurements on the days indicated in Figure 4a. Animal activity also was continuously monitored using an automated, digital platform in order to calculate the Digital Arthritis Index (DAI), an activity-dependent index of RA disease progression^35^. This index accounted for the baseline movement of the rats, which was calculated from the mobility of each animal during Days -2 to 5. An increase in the DAI corresponded with reduced mobility, which in turn correlated with increased disease severity.

From the ten candidates tested (Table S4), exenatide (10μg/kg), olopatadine (2mg/kg), and TXR-112 (0.5mg/kg) showed promise. All of these molecules significantly improved standard endpoints in the CIA model (Fig. 4, Table S5-S7). Exenatide, a glucagon-like peptide-1 agonist, and olopatadine, an antihistamine, are currently approved for the treatment of type 2 diabetes and allergic conjunctivitis, respectively. TXR-112 is still under investigation, and therefore, its name and mechanism will not be discussed further here. On Day 13 and Day 14, treatment with exenatide significantly reduced mean limb inflammation scores compared to vehicle-treated animals (Fig. 4b; p = 0.0177), and this trend was observed until the end of the study. Olopatadine treatment reduced limb inflammation at multiple time points; however, none of these effects were statistically significant (Fig. 4b). Ankle size also significantly decreased in animals treated with exenatide and olopatadine compared to their respective controls (Fig. 4c; p = 0.0305 and p < 0.0001, respectively). Treatment with exenatide (Supplementary Movie S1a) and olopatadine also significantly improve DAI, which indicated partial preservation of mobility (Fig. 4d). As expected, the inflammation score and ankle size did not increase significantly in animals treated with dexamethasone (Fig. 4b and 4c). In addition, no significant changes in mobility were observed in this group (Fig. 4c). These results demonstrated that treatment with exenatide and olopatadine significantly lowered the arthritis index compared to vehicle-treated controls, and partially preserved mobility during late stages of CIA progression (Fig. 4d).

Histopathological analysis was performed on ankles from animals treated with exenatide and olopatadine to corroborate the holistic readouts above. This analysis measured several parameters including inflammation, cartilage degradation, and bone resorption (Fig. 5a and 5b). Treatment with both drugs tended to lower histological scores compared to their respective vehicle-treated animals (Fig. 5c).

**Fig. 5.**
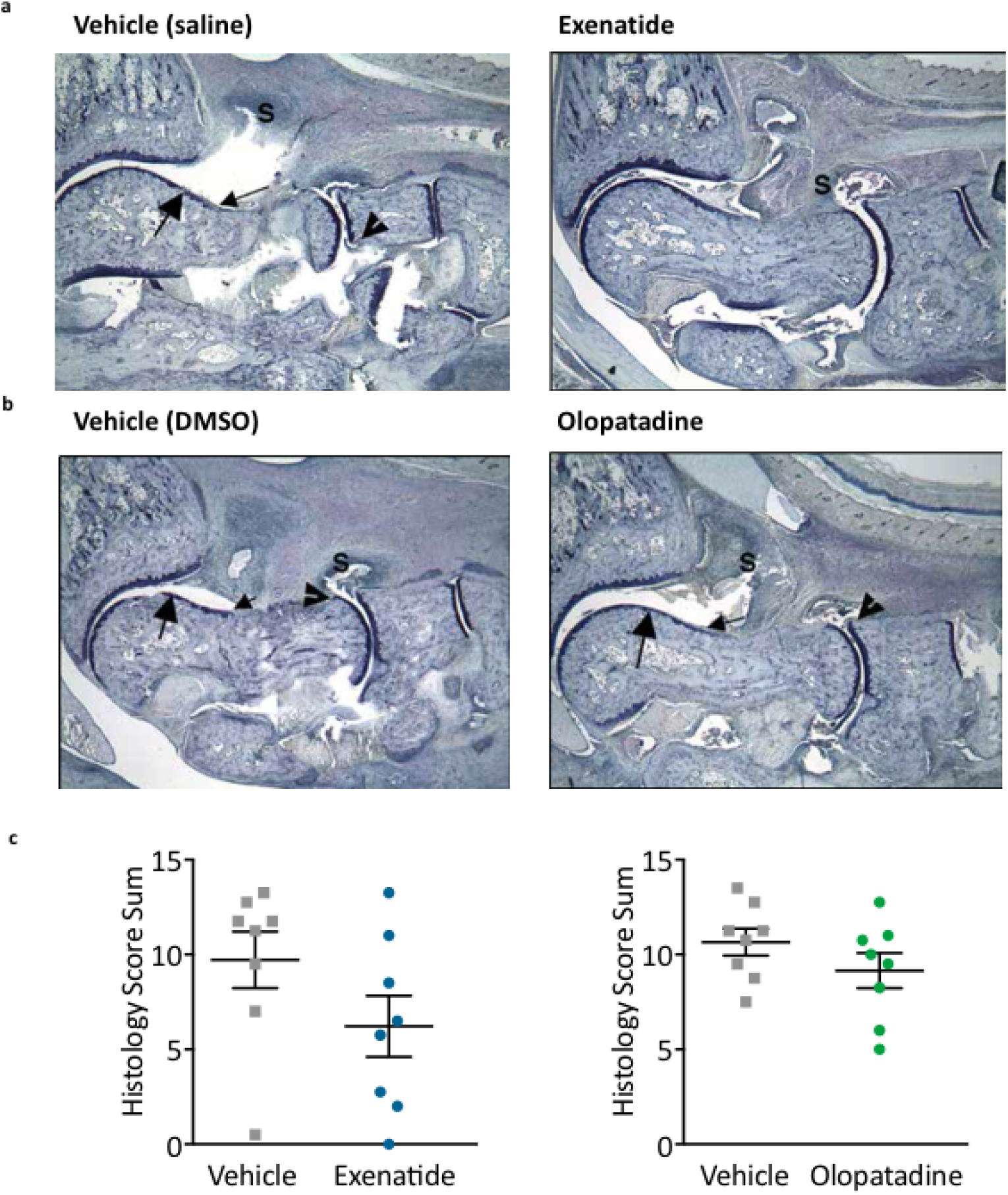
Histological assessment of exenatide and olopatadine treatment. (**a**) Micrographs of toluidine blue-stained ankle tissue obtained from exenatide- and vehicle-treated animals. (**b**) Micrographs of toluene blue-stained ankle tissue obtained from olopatadine- and vehicle-treated animals. (**c**) Histological sum score assessed from fixed ankle tissues. Data graphs represent the mean and SEM (n=8 per group). s: immune cell infiltrate; large arrow: cartilage; small arrow: pannus; arrowhead: bone resorption

In summary, our study demonstrated the power of using computational approaches to integrate large-scale, disparate data sources to identify novel and efficacious treatment of disease. The validation data from the *in vivo* preclinical model indicated that exenatide, olopatadine, and TXR-112 improved multiple outcomes of CIA, and showed therapeutic potential for the treatment of RA.

## Discussion

In this study, we rapidly identified high-potential treatments from among tens of thousands of possible drug candidates for RA using an integrative biomedical-informatics drug discovery platform. Identification of lead candidates occurred through three main phases over a 4-month period (Fig. 1). In Phase 1, 22,310 large and small molecules present in DrugBank^25^ and TTD^26^ were scored based on computationally-predicted therapeutic potential to yield 50 high-probability drugs candidates for RA. In Phase 2, algorithm evaluation and candidate due diligence identified 10 optimal candidates for further *in vivo* testing. In Phase 3, these 10 candidates were tested in preclinical studies and 3 lead candidates were identified.

The efficacy of our lead candidates, exenatide, olopatadine, and TXR-112, was demonstrated in a preclinical collagen-induced arthritis (CIA) model of RA. This is a highly reproducible model of severe polyarthritis that occurs when rodents develop immunity against collagen^36^. CIA in rodents phenocopies most of the features of RA: cartilage destruction, presence of immune cells in the joints, bone resorption, and inflammation. In particular, CIA in Lewis rats is a commonly used model for late, chronic stages of RA^37^. All three lead candidates improved limb inflammation, ankle size, and the Digital Arthritis Index (DAI) compared to their respective controls (Fig. 4, Fig. 5, Table S5-S7). Improvement in traditional endpoints, such as ankle joint measurements and clinical scores are indicative of reduced inflammation, erythema, and edema^32,38^. On the other hand, improvements in the DAI, a validated, activity-dependent, and indirect readout of disease severity, may indicate improved function and mobility^35^. Functional impairments, including activity and gait measurements have been described in RA models, and have been shown to better correlate with joint damage compared to clinical scores later in disease^39^. In RA patients, impaired mobility impacts productivity, participation in work and recreational activities, and quality of life ^2^; therefore, we included mobility as an efficacy outcome to capture the potential benefit of drug candidates in this key feature of RA. Even though the observed benefits of exenatide, olopatadine, and TXR-112 in the preclinical model are modest, we believe that our initial findings provide proof-of-concept, and further dose optimization and formulation of these drug candidates may yield greater efficacy. The doses chosen for this study were inferred from previous studies, where these drugs were investigated in the context of their canonical diseases in other organisms. Future studies should also involve replicating findings in other standard preclinical animal models of RA.

Current knowledge of these candidates’ targets and mechanisms for their original indication suggest that they do not act on common therapeutic targets for RA. We believe that these therapies improved *in vivo* endpoints by acting either on known targets in their approved indication, or engaging other uncharacterized therapeutic targets for RA. Elucidation of the precise mechanism of action of these drug candidates requires follow-up investigation as it was beyond the scope of this study. Future work will focus on identifying the target(s) that lead to improved outcomes in animal models of RA by leveraging what is already known about these candidates. As motioned previously, TXR-112 is a lead candidate that is still under investigation, therefore its name and mechanism will not be discussed further.

Exenatide is a glucagon-like peptide-1 (GLP-1) receptor agonist currently approved for the treatment of type 2 diabetes (T2D). It exerts its effects by increasing glucose-dependent insulin secretion, reducing glucagon levels, and promoting weight loss^40^. Links between RA and metabolic changes have been previously documented. Although the joints are proximally impacted in RA, cardiovascular disease, altered fat deposition, and insulin resistance are common co-morbidities^41^. Our gene expression meta-analysis also indicated an upregulation of insulin signaling in RA patients. The causal relationship between inflammation, RA, and insulin resistance is still unclear. At the molecular level, several reports suggest that metabolic symptoms associated with T2D could aggravate symptoms and disease progression in RA. In particular, hyperglycemia can increase plasma levels of TNF-α and IL-6^42^, both of which are pro-inflammatory cytokines with well-established roles in RA progression and as RA therapeutic targets^16^. Low doses of insulin can reduce levels of several factors implicated in immune cell migration and cartilage destruction in RA, including matrix metalloproteases and chemokines^43^. RA patients with the highest-grade inflammation also tend to be more insulin resistant than patients with low-grade inflammation^44^. These associations may point at a potential therapeutic benefit of repurposing T2D medications like exenatide in RA.

The specific anti-inflammatory and immunomodulatory properties of GLP-1 receptor agonists like exenatide are gaining recognition. Although these drugs treat diabetes by activating receptors expressed in the liver and pancreas, GLP-1 receptors are also expressed by various immune cells, including lymphocytes and macrophages^45–47^. Activity of GLP-1 receptor agonists on these cell types may augment therapeutic benefit in T2D, and may support the efficacy of these drugs in RA. Purified human GLP-1 was shown to inhibit chemokine-induced migration of lymphocytes^48^, and can regulate the proliferation of other immune cells^47^. Exenatide treatment *in vitro* can also shift macrophages from a pro-inflammatory M1 state to an anti-inflammatory M2 state^46^. Together, these studies suggest that GLP-1 receptor agonists like exenatide may indirectly modulate inflammatory and immune processes to relieve RA symptoms and slow disease progression.

Olopatadine is an anti-histamine and mast cell stabilizer indicated for allergic conjunctivitis and allergic rhinitis^49^. It works by antagonizing histamine H1 to inhibit the release of histamine from mast cells. Histamine is widely regarded as pro-inflammatory and induces chemotaxis of various other immune cell types that may potentiate inflammation and immune responses^50^. Rheumatoid synoviocytes express receptors for histamine H^51^, and administration of histamine to cultured synovial fibroblasts increased cell proliferation. Histamine can also stimulate expression of a matrix metalloprotease, an enzyme involved in cartilage degradation.

Olopatadine physically interacts with and likely has antagonistic activity towards several proteins in the S100 family of proteins^52^, which are implicated in a wide array of physiological processes, including inflammation and the immune response^53^. Specifically, olopatadine antagonizes the proinflammatory effects of S100A12, and inhibits the ability of this protein to induce chemotaxis for several immune cell types^49^. Several lines of evidence suggest that S100A12 protein levels correlate with RA disease activity or severity^54,55^. In addition, injecting S100A12 into mice mobilized neutrophils to sites of inflammation^56^. S100A12 may therefore reflect a novel therapeutic target for RA.

In addition to identifying new drug candidates and lead compounds, employing an unbiased computational approach such as ours to repurpose therapies can lead to a deeper understanding of RA pathophysiology. These drugs can be used to reveal previously unappreciated mechanisms involved in RA disease progression, which could in turn help identify new pathways and targets for drug development. In addition, identification of drugs that could be repurposed for RA can also serve as lead tool compounds which can be refined to produce novel candidates that can be more effective in treating RA. Results reported in this study validate our computational approach to drug discovery, and can easily be expanded to find novel mechanisms and therapies in other disease areas. This powerful approach does have limitations, the foremost of which is the requirement of sufficient quantities and diversity of quality data for the disease of interest and drugs investigated.

In summary, using a proprietary biomedical-informatics drug discovery platform, we processed patient-derived biological and clinical data along with data associated with over 22,000 existing drugs to identify candidates for drug repurposing in RA. Results in the CIA model suggest that exenatide, olopatadine, and TXR-112 improve endpoints by acting on either the known targets in their approved indication, or yet-to-be-discovered targets, that are being engaged in a way that is therapeutic for RA. This computational approach can be applied to numerous other disease indications to dramatically expedite drug discovery.

## Materials and Methods

### Gene expression meta-analysis data

#### Molecular data sources

All RA patient-derived gene expression datasets from Gene Expression Omnibus^57^ were reviewed for sustainability with our meta-analysis. We required there to be healthy controls as part of the experimental design and the datasets went through several quality control checks, including ArrayQualityMetrics™ ^58^. As a result, a total of 9 gene expression datasets from diverse disease-relevant tissues (synovial tissues, T-cell sub-types, and whole blood) were included (GSE20098, GSE57383, GSE57405, GSE45291, GSE7524, GSE1919, GSE55235, GSE12021, GSE55457, GSE10500, and GSE29746). Raw data were downloaded and normalized using RMA (R package Affy)^59^ when such data were avaiable. For all datasets, Limma ^60^ was used to identify significant differentially expressed genes. Default settings were used for all packages.

#### Drug-protein interaction network for rheumatoid arthritis

Scores for drug-protein associations were assigned as previously described^61^. Briefly, differentially expressed genes were mapped to proteins using UniProt identifiers. Differentially expressed proteins in RA, drugs linked by drug-protein interactions (DrugBank.ca v4.0^25^ and Therapeutic Target Database (TTD)^26^), and proteins linked by protein-protein interactions (PPI; Dr. PIAS^27^), were integrated in a network (Fig. 2). Drugs were scored according to: 1) the number of proteins differentially expressed in RA with which that drug’s targets interact, 2) the confidence and directionality of those interactions, and 3) the consistency of differential protein expression across individual RA datasets.

#### Enrichment of existing RA treatments among drug predictions

A method similar to Gene Set Enrichment Analysis (GSEA)^62^ was used to determine the enrichment of existing treatments among the highest scoring drugs identified using any given method as previously described^61^.

#### Drug-disease co-occurrence and aggregate scoring

All event records from the first quarter of 2013 through the third quarter of 2014 were downloaded from the FDA Adverse Event Reporting System (FAERS). All drug and disease indications were extracted from each event; here an event is a single reporting and thus consists of a single patient. Drugs were identified using the name listed in FAERS and matching that to the canonical names or synonyms provided by DrugBank or TTD. If perfect matches were not present, drug names were split on parentheses (often used to denote dosage), and the resulting name used to match to DrugBank and TTD. Unmappable drug names were excluded from subsequent analysis.

To calculate the strength of association between each drug and rheumatoid arthritis, Fisher’s exact test was used to calculate the odds ratio of taking that drug and having RA along with the corresponding two-tailed p-value, which was multiple hypothesis corrected using Benjamini-Hochberg. The odds ratio for each drug was converted to a score by taking the absolute value of the log2 of that ratio. Drugs with a corrected p-value greater than 0.05 had their scores set to 0, in effect removing them from the analysis.

Scores derived from each of the above methods were integrated using our proprietary machine learning algorithm.

### Candidate review

Following prioritization of drug candidates using twoXAR’s proprietary algorithms, the top 50 candidates were manually inspected to ensure novelty and applicability for the project. Any candidates previously researched or patented for treatment of RA were excluded. In addition, we required all drugs to be commercially available and eliminated any candidates with unacceptable toxicity profiles for chronic use.

### Collagen-induced arthritis (CIA) Induction

Female Lewis rats between 7–10 weeks of age and weighing approximately 175g were obtained from Charles River (Wilmington, MA) and habituated for 7 days prior to CIA induction. Rats were housed singly in 12-hour light/dark cycles with unrestricted access to food and water. During this habituation period, rat mobility and weight were recorded and used to randomly assign animals to treatment groups. To prepare the inoculum for CIA induction, lyophilized porcine type II collagen obtained from Chondrex (Redmond, WA) was added to 0.01N acetic acid to a final concentration of 2 mg/mL. The collagen inoculum was then emulsified with an equal volume of incomplete Freud’s adjuvant (IFA) obtained from Sigma-Aldrich (St. Louis, MO). Rats were then anesthetized with 5% isofluorane, the base of the tail was shaved, and 200 μL of inoculum was injected at two sites at the base of the tail. This marked Day 0 of the study. A second, single booster injection of collagen-IFA (100 μL) was administered at the base of the tail 7 days after initial inoculation (Day 7), as previously described^63,64^. All animal experiments were conducted at Vium, Inc. (San Mateo, CA) and the animal use protocol was approved by Vium’s Institutional Animal Care and Use Committee (IACUC).

### Candidate Drug and Control Dosing

Each drug candidate was tested using three groups consisting of 8 rats each. The number of rats included in each group was determined by power analysis. Three testing doses (high, mid, and low) for each candidate were determined by identifying the maximally tolerated and minimally efficacious doses used in previous non-RA studies in rats. If data in rats were not available, doses were inferred based on studies in other organisms. Rats were randomly assigned to treatment groups according to data on mobility and body weight collected during the habituation period. In Lewis rats, arthritis symptoms develop approximately 10 days after immunization with collagen and IFA^29^. Starting on Day 9, animals were treated daily with one of three doses of each drug candidate or the drug vehicle as indicated in Table S8. TXR-112 (Sigma; St. Louis, MO) and exenatide (Cayman Chemical; Ann Arbor, MI) were formulated in saline. Olopatidine (Cayman Chemical; Ann Arbor, MI) was formulated in 1% DMSO. Dexamethasone (MWI; Grand Prairie, TX) was formulated in 0.5% methylcellulose and administered orally. An appropriate route of administration (intraperitoneal, subcutaneous, or oral) was determined for each drug candidate (Table S8). All dosing was performed by personnel blinded to the treatment group. Treatment continued until Day 16. All animals were euthanized by CO_2_ inhalation on Day 17 of the study. The right and left ankles were removed and placed in 10% neutral buffered formalin for histological analysis.

### Efficacy Outcomes

Several standard outcomes were measured to examine the severity of arthritis development and efficacy of candidate and control treatments ^63,65^. Both ankle joints were measured by caliper prior to CIA induction (day 0) and again on days 7 and 17 for all drugs. For animals treated with olopatidine only, an additional ankle measurement was collected on day 14. Ankle sizes for both right and left were averaged during analysis and normalized to the mean size for each group at Day 0. On days 0, 7, and 9–16, animals were scored for hind limb inflammation as described in Table 1. Scores for both limbs were summed during analysis. In addition to standard measures, the validated Digital Arthritis Index (DAI) was continuously collected as previously described^35^. Briefly, for each individual animal, salient features of motion, including maximum speed during the dark cycle, was extracted and aggregated by study day, then normalized to a baseline period (Study Days -2 to 5, excluding induction Day 0). An increased DAI score corresponds to reduced mobility relative to baseline.

**Table 1.** Hind limb inflammation scoring. Scores for hind limb inflammation were assigned based on the criteria described.

| Score | Limb inflammation assessment |
| --- | --- |
| 0 | Normal |
| 1 | Mild, but definite redness and swelling of the ankle, or apparent redness and swelling limited to individual digits, regardless of the number of affected digits |
| 2 | Moderate redness and swelling of the ankle |
| 3 | Severe redness and swelling of the entire paw including digits |
| 4 | Maximally inflamed limb with involvement of multiple joints |

### Histopathology and microscopic damage assessment

Formalin-fixed ankle joints were submitted to Bolder BioPATH (Boulder, CO) for processing, sectioning, and evaluation. Decalcified ankles were cut in half longitudinally and the two halves were embedded together in a paraffin block. Sections were cut from each block and stained with toluidine blue. Joints were examined microscopically in a blinded fashion by a board certified veterinary pathologist and observations were entered into a computer-assisted data retrieval system. Ankle inflammation, pannus, cartilage damage, bone resorption, and new periosteal bone formation were scored and the summed score was calculated for each ankle.

## List of Supplementary Materials

**Table S1.** Drugs in clinical trials for RA.

**Table S2.** Reactome pathway analysis of differentially expressed proteins in rheumatoid arthritis samples versus controls.

**Table S3.** Top 100 candidates identified using the aggregated scoring method.

**Table S4.** List of 10 drug candidates selected for *in vivo* testing in cartilage-induced arthritis animal model.

**Table S5.** Raw inflammation scores of animals treated with exenatide, olopatidine, and TXR-112.

**Table S6.** Raw ankle measurements of animals treated with exenatide, olopatidine, and TXR-112.

**Table S7.** Raw Digital Arthritis Index Scores of animals treated with exenatide, olopatidine, and TXR-112.

**Table S8.** Route of administration and dosage used for exenatide, olopatidine, and TXR-112 in *in vivo* studies.

Movie S1. Digital recording of exenatide (b)- and vehicle (a)-treated animals.

## Acknowledgments

The authors thank D. Hutto and T. Heuer for suggestions to the manuscript.

## Author contributions

A.C.D., C.F., and A.A.R. designed and built the integrative bioinformatic platform with advice and feedback from M.S.. A.C.D. and E.S.N. loaded and annotated the disease-specific data. A.C.D., I.H., E.S.N., A.M.R., M.S.C., M.S., and A.A.R. reviewed and selected final drug candidates. I.H. designed the *in vivo* validation experiments that were directed and carried out by M.R. and L.S.. L.S., D.F., G.F., I.H., and E.S.N. analyzed the *in vivo* results. A.C.D and S.M. wrote the paper with help and feedback from E.S.N., A.M.R., and M.S..

